# Microbial trend analysis for common dynamic trend, group comparison and classification in longitudinal microbiome study

**DOI:** 10.1101/2020.01.30.926824

**Authors:** Chan Wang, Jiyuan Hu, Martin J. Blaser, Huilin Li

## Abstract

**Motivation:** The human microbiome is inherently dynamic and its dynamic nature plays a critical role in maintaining health and driving disease. With an increasing number of longitudinal microbiome studies, scientists are eager to learn the comprehensive characterization of microbial dynamics and their implications to the health and disease-related phenotypes. However, due to the challenging structure of longitudinal microbiome data, few analytic methods are available to characterize the microbial dynamics over time.

**Results:** We propose a microbial trend analysis (MTA) framework for the high-dimensional and phylogenetically-based longitudinal microbiome data. In particular, MTA can perform three tasks: 1) capture the common microbial dynamic trends for a group of subjects on the community level and identify the dominant taxa; 2) examine whether or not the microbial overall dynamic trends are significantly different in groups; 3) classify an individual subject based on its longitudinal microbial profiling. Our extensive simulations demonstrate that the proposed MTA framework is robust and powerful in hypothesis testing, taxon identification, and subject classification. Our real data analyses further illustrate the utility of MTA through a longitudinal study in mice.

**Conclusions:** The proposed MTA framework is an attractive and effective tool in investigating dynamic microbial pattern from longitudinal microbiome studies.

## Background

The human microbiota represents a complex and rich ecosystem of over 100 trillion microbial cells, playing a fundamental role in maintaining health and driving disease [1, 2]. Considering that the human microbiota is inherently dynamic and can be substantially altered by many factors at any time point either temporally or permanently, recent microbiome studies [2–8] have shifted their research design from cross-sectional or case-control studies to longitudinal analyses. Rather than untangling the associations between microbes and a wide range of diseases at a fixed time point [2, 9–13], longitudinal microbiome studies target understanding how the dynamic changes in microbiome are linked with disease susceptibility. For example, the Integrative Human Microbiome Project [2, 3] was designed to comprehensively characterize the dynamic changes in human microbiome in three disease-specific cohorts: pregnancy and preterm birth, onset of IBD, and onset of type 2 diabetes. Serrano et al. (2019) [5] found that the vaginal microbiome shifted during pregnancy toward a more homogeneous *Lactobacillus*-dominated profile which is associated with the risk of preterm birth. Lloyd-Price et al. (2019) [6] reported that periods of IBD disease activity were distinguished by gut microbiome stability, with taxonomic, functional, and biochemical shifts of microbiota. In studies related to diabetes, Zhou et al. (2019) [4] showed that healthy profiles displayed diverse patterns of intra- and inter-personal variability, and that different microbial profiles were identified between insulin-resistant participants and insulin-sensitive participants at baseline and during stress responses. Such scientific results provide insights into the characterization of the microbial dynamics and raise further questions about understanding these underlying microbial dynamics as well. Do the microbial dynamics significantly contribute to the group differentials? If so, which specific microbes dominate them? Can subjects be classified based on their microbial dynamics?

To address such questions, recent efforts have focused on analyzing the longitudinal microbiome data. Several parametric methods have been proposed to elucidate the microbial dynamic changes, including mixed-effects model [14, 15], generalized Lotka-Volterra equations [16, 17] and time series models [18–20]. While those methods provide capabilities to capture the microbial dynamics and identify the time-dependent taxa, they assume that the abundance of an individual taxon changes either at a fixed rate or in an autoregressive pattern, which cannot always be justified. In contrast, a nonparametric approach (Permuspliner) uses the loess spline to examine whether one taxon’s changing pattern over time is significantly different between two groups or not [21]. However, the number of taxa that need to be tested can inevitably affect the statistical power after the multiple testing corrections due to the high dimensionality of the microbiome, even at high taxonomic ranks.

Given that most available methods to analyze longtitudinal microbiome data focus on modelling the trends of a single taxon, while microbiota often acts as an integrated community, we propose a microbial trend analysis (MTA) which can capture the overall community level microbial dynamic pattern for a group of subjects. In particular, following the path of the principal trend analysis (PTA) [22], MTA integrates the spline-based method for time-course data analysis and principal component analysis for dimension reduction to extract the dynamic patterns from a group of subjects. Matrix decomposition and lasso technique are used to address the high-dimensionality feature, and the graph Laplacian penalty [23–26] is used to incorporate phylogenetic tree structure into analysis. In combination, these help MTA to identify the dominant taxa which contribute to the common trends simultaneously. Based on the proposed MTA, if the differential trend is significant, we further propose a microbial trend group differential test which can identify the contributing taxa, and a distance-based classification algorithm to assign a group label for a given subject, which is potentially useful for considerations of personalized treatments.

The remainder of this paper is organized as follows. Firstly, we introduce in detail the three components of the proposed MTA framework: MTA model, group differential test, and classification algorithm. Secondly, we conduct extensive simulations to evaluate MTA in testing and classification. Thirdly, we illustrate the utility of MTA to investigate the relationship between antibiotic usage and the microbial dynamics using a longitudinal murine study. Finally, we conclude with an overall discussion.

## Methods

### Microbial trend analysis

Suppose there are *N* subjects in a given group. Let *Y*_*npt*_ be the relative abundance of the *p*^th^ taxon of subject *n* at time point *t, p* = 1, …, *P*, and *t* = 1, …, *T*, and 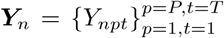 be a *P* × *T* matrix of the relative abundances of subject *n*. To extract the common trends shared by all *N* subjects, we consider the following optimization problem, as illustrated in PTA [22],

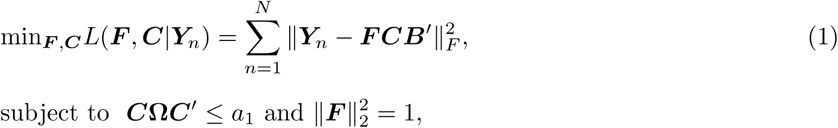

where ***F*** = (***f***_1_, …, ***f***_*M*_) is a *P* × *M* matrix of factor scores, ***C*** = (***c***_1_, …, ***c***_*M*_)′ is an *M* ×(*T* +2) matrix of spline coefficients, 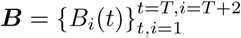 is a *T* ×(*T* +2) matrix containing the cubic spline basis and ‖·‖_*F*_ is the Frobenius norm. Denote ***Q*** = ***CB***′, which is an *M* × *T* matrix presenting the top *M* common trends across *T* time points. 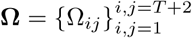 is a (*T* + 2) × (*T* + 2) matrix with 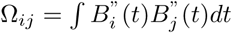, *i, j* = 1, …, *T* + 2. *M* can be either predetermined by the user or determined by specifying the percentage of variance to be explained by the top common trends (see Algorithm 1 for details). The tuning parameter *a*_1_(≥ 0) controls the smoothness of trends with a smaller value of *a*_1_ producing smoother trends. Note that ***F*** and ***C*** are identical for all subjects respectively, for they integrate information from all subjects and represent the common trends ***Q*** shared by all subjects. The elements of ***f***_*m*_ are the weights of *P* taxa on the *m*^th^ common trend ***Bc***_*m*_, of which a higher absolute value indicates a stronger contribution to this common trend. With this modeling, we can find the top *M* common trends from a group of subjects, and quantify the contributions of *P* taxa to each of the common trends.

In the microbiome data, the number of taxa *P* is usually far larger than the sample size *N* or the number of time points *T*, and it is reasonable to assume that only a few taxa make substantial contributions to the community level dynamic trends [27]. Thus, following PTA, we impose Lasso penalty on the factor score parameter ***F*** to ease the high-dimentionality problem and identify these key taxa. Another important feature of microbiome data is the phylogenetic tree structure that describes the evolutionary relationships among taxa. Evolutionarily related taxa tend to have similar effects on human phenotypes or diseases [23, 24, 26, 28]. In the existing studies, incorporating the phylogenetic tree information into the analysis improves both statistical power and biological interpretability [23–26]. To take this factor into account, we integrate the following Laplacian penalty in the optimization problem (1),

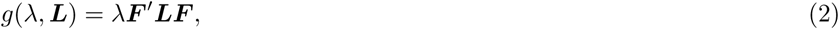

where *λ*(≥ 0) is the tuning parameter. The Laplacian matrix ***L*** is determined by the phylogenetic tree and constructed using a similar approach to that described in Chen et al. (2012) [23]. Detailed construction of ***L*** is given in the Additional file 1: Section S1. With these two additional penalties, we introduce two Lagrange multipliers to rewrite the optimization problem (1) as

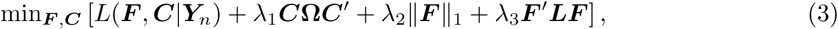

subject to 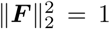, where *λ*_1_(≥ 0), *λ*_2_(≥ 0) and *λ*_3_(≥ 0) are tuning parameters that control smoothness of the common trends, sparsity, and smoothness of the estimated factor scores based on the phylogenetic tree distance, respectively. As demonstrated in Zhang and Davis (2013) [22], the optimization problem (3) is biconvex, though not convex. Thus we adapt an iterative nonlinear partial least square algorithm in Algorithm 1 to get estimates 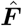 and ***Ĉ*** by minimizing the optimization problem (3) [22, 29]. The detailed derivation is provided in the Additional file 1: Section S2.

#### Algorithm 1: Nonlinear iterative algorithm for MTA

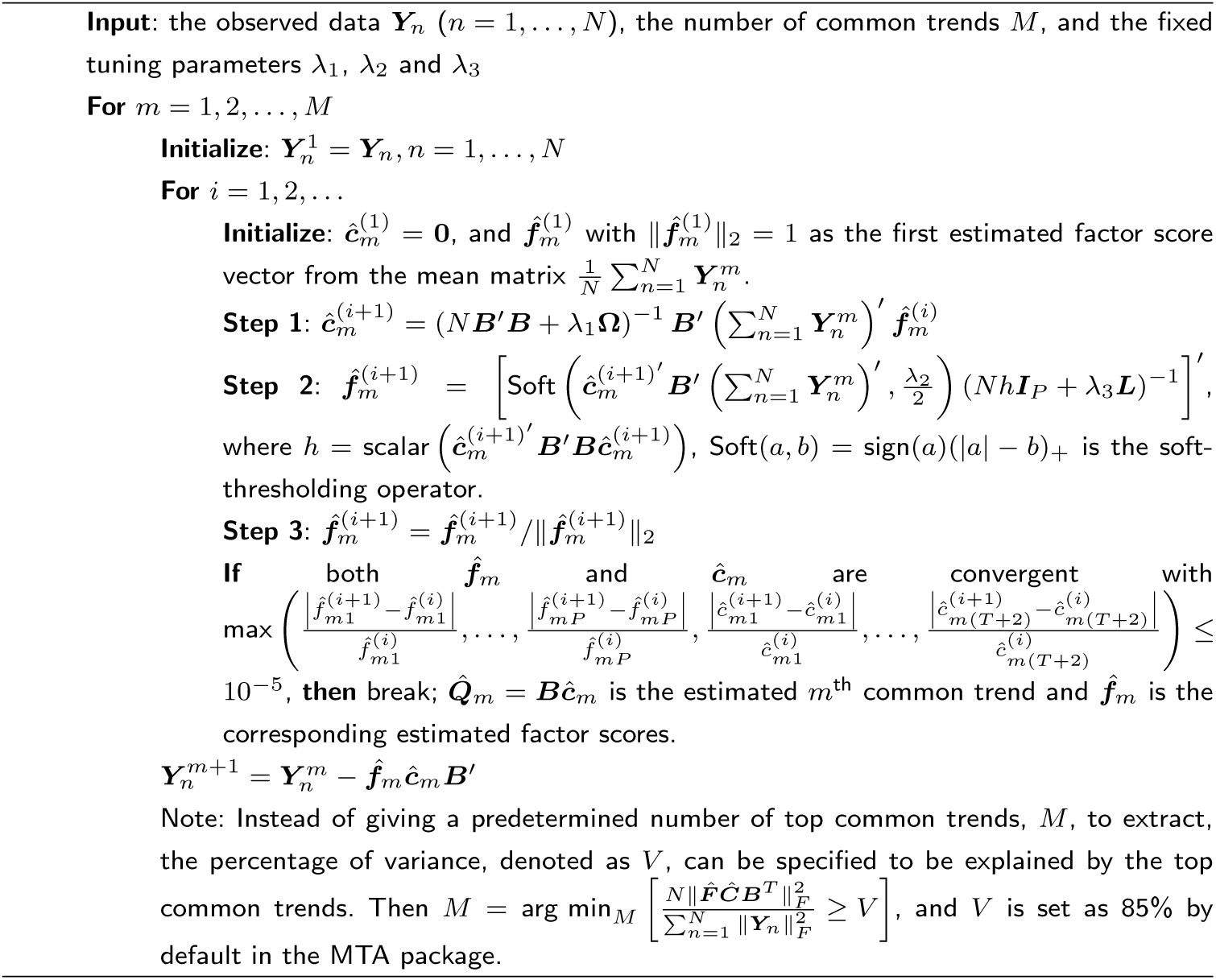

Given the observations ***Y***_*n*_, *n* = 1, …, *N*, the number of top common trends *M* or the percentage of variance *V* for the top common trends to explain, and the tuning parameters *λ*_1_, *λ*_2_ and *λ*_3_, Algorithm 1 solves for the estimated top common trends 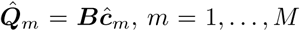, shared by all *N* subjects, which are the linear combinations of the relative abundances of all *P* taxa across *T* time points, and 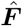 indicates the dominant taxa and their contributions to the common trends.

Note that the choices of tuning parameters in Algorithm 1 are crucial for extracting the common trends and we use the *K*-fold cross-validation (CV) [22, 23, 30] to determine the values of *λ*_1_, *λ*_2_ and *λ*_3_. Specifically, we randomly split the indexes of all subjects 1, …, *N* into *K* exclusive folds with roughly equal sample size *G*_1_, …, *G*_*K*_, and take the subjects ***Y***_*n*_, *n* ∉*G*_*k*_, as the training set and the remaining subjects as the testing set in the *k*th cross-validation, *k* = 1, …, *K*. With the given candidate values of the tuning parameters ***λ*** = (*λ*_1_, *λ*_2_, *λ*_3_), we obtain estimates 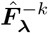 and 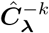 from the training set with Algorithm 1 in the *k*th cross-validation. The predicted group-level microbial composition for this training set therefore is 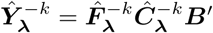. Then the total squared error on the testing set ***Y***_*n*_, *n* ∈ *G*_*k*_, is defined as 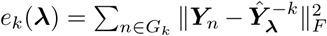, and the average mean squared error over all *K* folds is recorded as

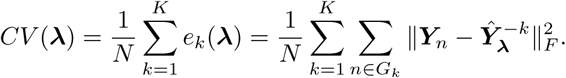

Finally, we select ***λ***\* = arg min_***λ***_*CV* (***λ***) as the optimal tuning parameters.

### Group comparison

In the subsection above, we proposed the MTA method to capture the common dynamic trends shared by all subjects in a given group and identify the dominant taxa contributing to these common trends. We next apply the MTA method to evaluate how different two groups are in terms of their microbial dynamic pattern, and if the group difference is significant, quantify the differentiating taxa. The proposed group test is based on a difference array constructed using pseudo samples and permutation technique. Specifically, let 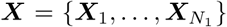 and 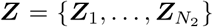 be the arrays of relative abundances for *N*_1_ controls and *N*_2_ cases with *P* taxa across *T* time points (here, ***X***_*n*_ and ***Z***_*n*_ have a similar definition as ***Y***_*n*_ in the subsection above), respectively. The difference array then is constructed as

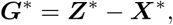

where 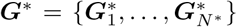 and 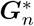 represents the relative abundance difference between randomly paired subjects from case and control groups with 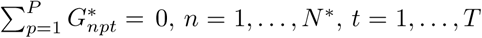, and *N*\* = min (*N*_1_, *N*_2_). 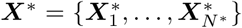 and 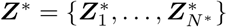 are the permuted controls and cases respectively.

Under the null hypothesis of no difference between cases and controls, the expectation of 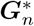 should be a *P* × *T* zero matrix and the difference array ***G***\* thus has only noise signal. On the other hand, under the alternative hypothesis, that control and case groups hold different dynamic patterns, 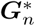 is expected to be time-related and to contain some common trends illustrating the underlying differences between cases and controls, which should differ from the random noise signal under the null. Thus, we propose a test to evaluate the dynamic differences between cases and controls by comparing the trends extracted from ***G***\* to the noise trend which is randomly arranged around 0 across all time points. Specifically, the test is evaluated based on the resampling (or bootstrapping) method [31], with details described in Algorithm 2.

#### Algorithm 2: Hypothesis testing for group comparison

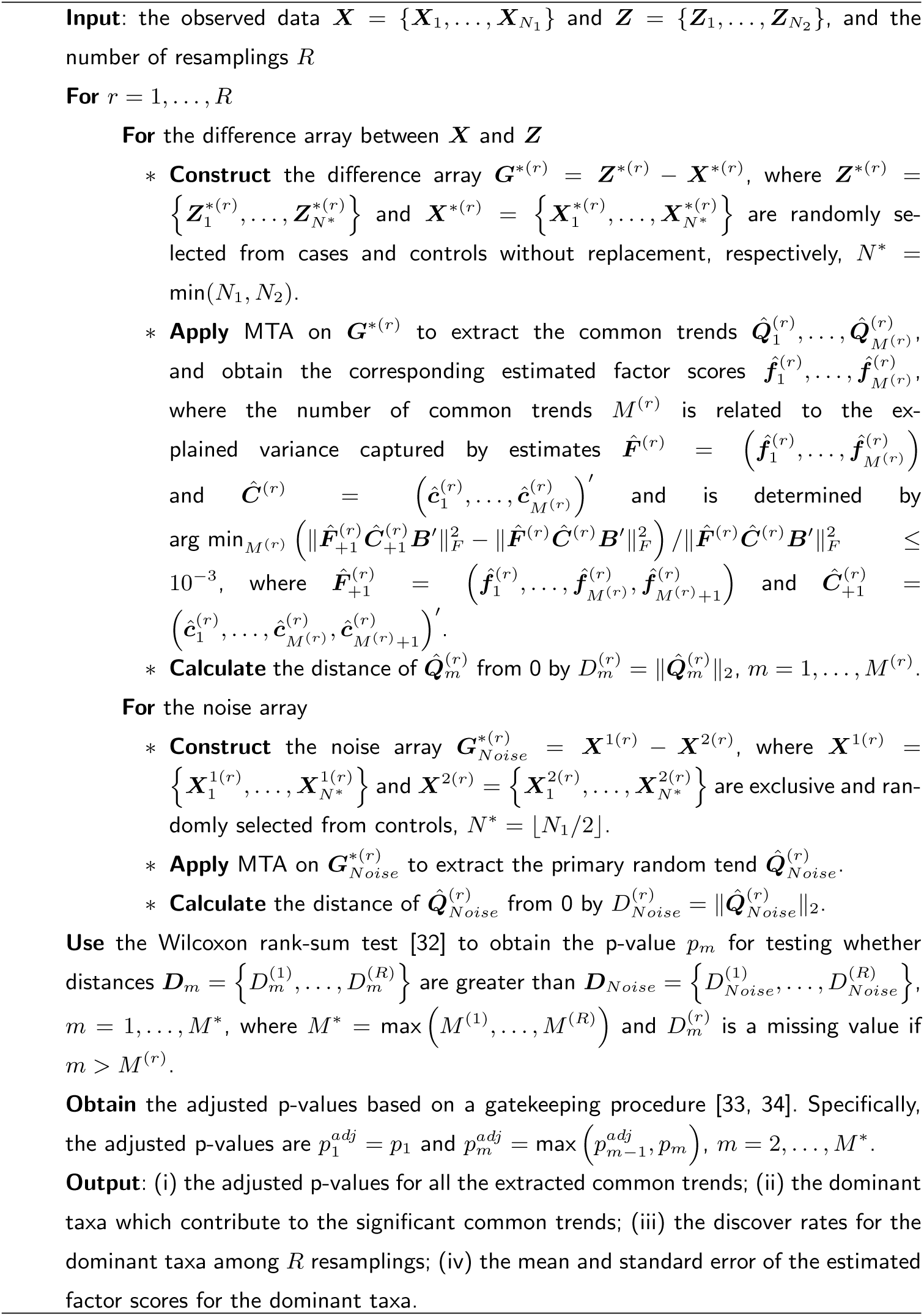

Note that Algorithm 2 consists of several components used in constructing the hypothesis test for assessing the microbial dynamic group differences. Firstly, we determine the number of common trends extracted from the difference array ***G***\*^(*r*)^, *M* ^(*r*)^, by the criterion that the increasing rate of the explained variance is equal to or less than 10^−3^, to capture most of the information shared by all pseudo subjects. Secondly, we employ the straightforward and widely used Euclidean distance to measure the differences between the common trends extracted from ***G***\* and the random trend from 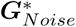. Thirdly, we use the Wilcoxon rank-sum test to check whether distances ***D***_*m*_ are greater than ***D***_*Noise*_, and provide the corresponding p-value *p*_*m*_ for the *m*^th^ common trend, *m* = 1, …, *M* *. Finally, multiple testing correction is needed to adjust these individual p-values to control the overall error rate. Since these *M* * hypothesis tests exhibit a hierarchical structure in which the information captured by the common trend ***Q***_*m*_ monotonically decreases as *m* increases, we use a gatekeeping procedure [33, 34] to take care of this hierarchical structure and obtain the adjusted p-values 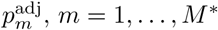. This implies that the *m*^th^ common trend is meaningful only if the previous one has a significant difference.

### Classification

In addition to the common dynamic trends and the estimated factor scores, the MTA method provides the predicted microbial composition matrix: 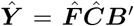 at the group level. Since the predicted microbial composition matrix integrates information from all subjects of a given group and represents the unique and underlying pattern defined by this group, it can serve as a prototype of this group. Therefore, we propose a distance-based classification algorithm (Algorithm 3) to classify subjects based on their distances from the predicted microbial matrices of two different groups in the training set. Specifically, suppose there is a training set involving controls ***X*** and cases ***Z***, and a new subject ***Y***_new_ with an unknown group label. Firstly, we apply the MTA method to controls ***X*** and cases ***Z*** separately to obtain their corresponding predicted matrices 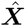 and 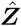. Here the number of common trends *M* integrating the predicted microbial matrix is determined by the proportion of explained variance, as illustrated in Algorithm 1. Secondly, we calculate distances of the new subject ***Y***_new_ from 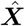 and 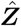, which are defined as 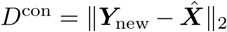 and 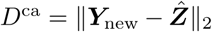 respectively. Finally, this new subject ***Y***_new_ is classified as a case if *D*^con^ > *D*^ca^, otherwise as a control.

#### Algorithm 3: Classification

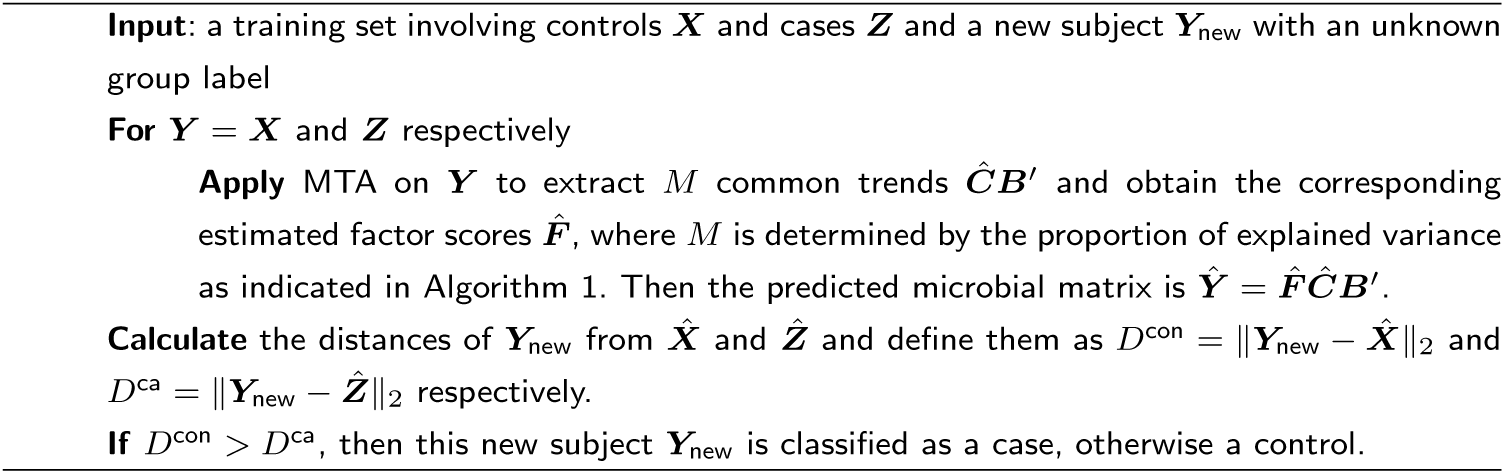

## Simulation study

We constructed extensive simulations to evaluate the performance of the proposed MTA method in both group comparison (Algorithm 2) and classification (Algorithm 3). Specifically, the performance of group comparison is evaluated in terms of the empirical type I error rate and statistical power at the group level, as well as in terms of sensitivity and specificity at the taxon level, compared to the competing method Permuspliner which has been the only available method proposed to evaluate microbial dynamic patterns between groups so far (R package: Shields-Cutler et al., 2018 [21]). Permuspliner tests the differences in microbial abundance between two groups over the longitudinal time course based on the loess splines [35, 36]. We evaluated the performance of classification in terms of receiver operating characteristic (ROC) curve and area under the curve (AUC) [37].

### Simulation design

Following Wang et al. (2019) [38], we designed our simulation at the genus level based on the real data illustrated in the “Real data analysis” section [39]. The real data include 106 genera for 20 control and 17 sub-therapeutic antibiotic treatment (STAT) male mice across 4 time points (weeks 3, 6, 10 and 13). After filtering out those genera that appear in < 10% of samples or with mean proportions < 10^−4^, 35 taxa were included in the analysis. We simulated the microbial relative abundances at each time point for cases and controls respectively from the Dirichlet distribution. Specifically, the mean relative abundance of the *p*^th^ taxon for a subject at the *t*^th^ time point *O*_*pt*_ is assigned as below for *p* = 1, …, *P* and *t* = 1, …, *T*,

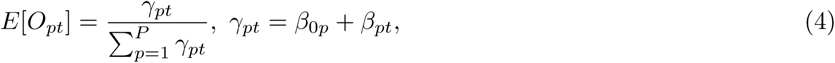

where ***β***_0_ = (*β*_01_, …, *β*_0*P*_)′ represents the baseline relative abundances for *P* = 35 taxa, which were set as the corresponding estimates from the control mice using R package dirmult [40]. *β*_*pt*_ is the deviation from the baseline relative abundance 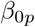 of the *p*^th^ taxon at the *t*^th^ time point, which represents the group difference.

To estimate the empirical type I error rate, the relative abundances of controls and cases were generated under the null hypothesis of no group difference by setting *β*_*pt*_ = 0, *p* = 1, …, *P* and *t* = 1, …, *T*.

To evaluate the statistical power, we designed three scenarios with two different common trends between control and case groups, of which 3 dominant taxa (about 10% of the total) contributed to the first one and 2 dominant taxa (about 5%) contributed to the second one. Denote the set of the indices of these 5 dominant taxa as ∧_1_ and ∧_2_ and their corresponding deviations as 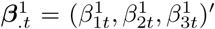 and 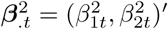. Following Zhang and Davis (2013) [22], they were assigned as

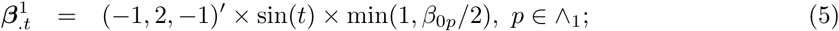

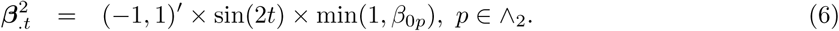

Furthermore, the ∧_1_ and ∧_2_ were determined while taking the phylogenetic tree structure into account as follows. We firstly partitioned the 35 taxa into 5 clusters using the partition around medoids algorithm [41] on the patristic distance in the real phylogenetic tree. Secondly, we randomly selected two clusters and then randomly selected 3 and 2 taxa from these two clusters without replacement as the taxa in ∧_1_ and ∧_2_, respectively. Consequently, the dominant taxa that contributed to the group differences were phylogenetically related.

Three different simulation scenarios were constructed to thoroughly illustrate the performance of the proposed MTA method and the competing method Permuspliner: (1) the relative abundances in the control group were generated with *β*_*pt*_ = 0, while the relative abundances in the case group were generated with 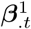 for the taxa in ∧_1_ and *β*_*pt*_ = 0 for *p* ∉∧_1_; (2) the relative abundances in the control group were generated with *β*_*pt*_ = 0, while the relative abundances in the case group were generated with 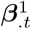 and 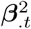 for the taxa in ∧_1_ and ∧_2_ respectively and *β*_*pt*_ = 0 for *p* ∉∧_1_ ∪ ∧_2_; (3) the relative abundances in the control group were generated with 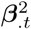 for the taxa in ∧_2_ and *β*_*pt*_ = 0 for *p* ∉∧_2_, while the relative abundances in the case group were generated with 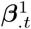 and 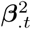 for the taxa in ∧_1_ and ∧_2_, respectively, and *β*_*pt*_ = 0, *p* ∉∧_1_ ∪ ∧_2_; *p* = 1, …, *P, t* = 1, …, *T*. Hence, there is only one different common trend between control and case groups in both scenario 1 and scenario 3, but two different common trends in scenario 2. Finally, we considered two equal sample sizes *N*_*case*_ = *N*_*control*_ = *N* = 20 or 30, and two numbers of time points *T* = 10 or 20.

### Simulation results for group comparison

Since the common trends extracted by Algorithm 2 manifest the difference between case and control groups and have a hierarchical structure as well, the overall empirical type I error rate and power for the proposed MTA method were defined as the proportion of the adjusted p-values for the first common trend less the given significance level (usually 5%) with 1000 independent replications under the null and alternative hypotheses respectively. Since the competing method Permuspliner works at the taxon level and only provides the individual p-value for each taxon separately, its overall type I error rate and power were calculated as the proportion of at least one taxon that had a significant adjusted p-value with the Benjamini-Hochberg (BH) correction for multiple comparisons [42]. The number of resamplings in Algorithm 2 was set as *R* = 50.

#### Type I error rate and power

The first group of bars in each subfigure of Figure 1 reports the empirical type I error rates of the proposed MTA method and the competing method Permuspliner. They are all around the nominal significance level 5%, except that when sample size *N* = 20 and the number of time points *T* = 20, both methods, especially Permuspliner, have slightly conservative type I error rates. Thus, both are statistically valid tests to differenciate microbial dynamic patterns between groups in the longitudinal study. The groups 2-4 in each subfigure of Figure 1 report the estimated powers of both methods with sample size *N* = 20, 30 and the number of time points *T* = 10, 20 in scenarios 1-3, respectively. The relative performance of these two methods is similar across scenarios 1-3 in all combinations of *N* and *T*. MTA is more powerful than Permuspliner in most of the scenarios, especially when the number of time points is large at *T* = 20. In addition, the power of MTA increases as *N* or *T* increases. Taking scenario 2 as an example, the estimated power of MTA increases from 37.1% with *N* = 20 and *T* = 10 to 66.0% with *N* = 30 and *T* = 10, and to 90.5% with *N* = 30 and *T* = 20. The number of time points *T* has an effect on the power of the competing method Permuspliner, since increasing time points from 10 to 20 diminishes its power dramatically in all scenarios. This is because Permuspliner is based on the area under the microbial splines across all time points to evaluate dynamic patterns. The area under the curve method is most effective when the temporal trend difference between the compared groups is relatively consistent along time, i.e. one trajectory is always above the other. However, when the microbial trends from two groups cross over, which is more likely to happen when the observed time period is longer, the area under the curve difference can even out and accordingly be reduced in power for differentiating dynamic patterns. Since in practice, microbial dynamic patterns are nonlinear and the group changing trends are more likely to intersect, as supported by our real data in Figure 5, the proposed MTA method has superior power.

**Figure 1:**
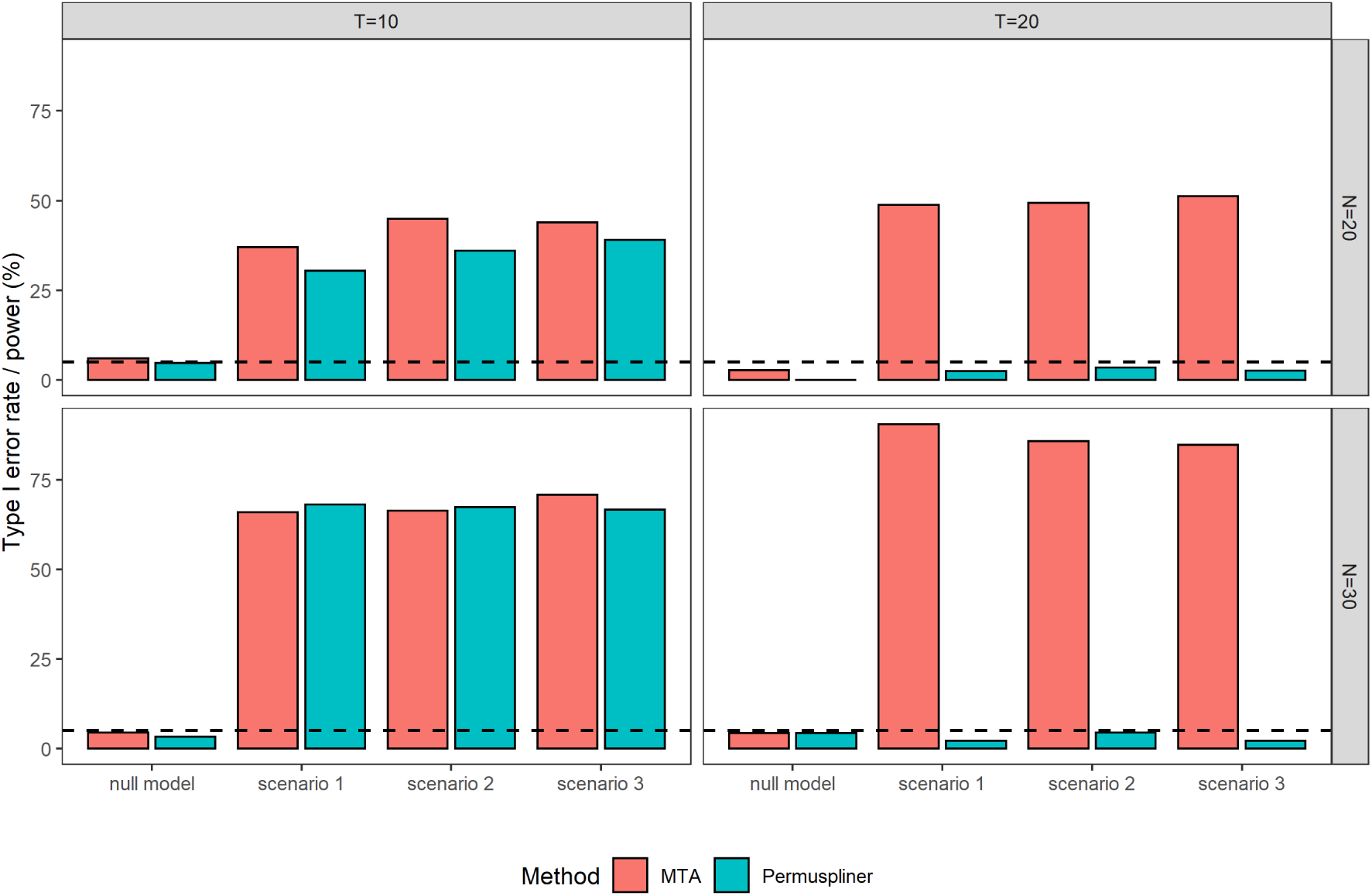
Empirical type I error rate and power (%) for testing the difference between case and control groups with sample size *N* = 20, 30 and the number of time points *T* = 10, 20 in null model and scenarios 1-3, respectively. The dashed line represents the given significance level 5%.

#### Sensitivity and specificity

The estimated powers above illustrate that the proposed MTA method has superior performance in testing the differences between two groups than the competing method Permuspliner at the group level. We further evaluate the performance of both methods at the taxon level in terms of sensitivity and specificity. Specifically, sensitivity is defined as the proportion of taxa correctly identified from 5 dominant taxa that truly contribute to the difference between two groups, and specificity is the proportion of uninvolved taxa that are correctly identified as such. Furthermore, the proposed MTA method determines whether a taxon was significantly contributing or not by the estimated confidence interval of its factor score with the number of resamplings *R* = 50 in Algorithm 2. For the competing method Permuspliner, the adjusted p-value with BH multiple comparison correction determines whether a taxon is significantly different between two groups or not.

Figure 2 presents the estimated sensitivity and specificity of the MTA and Permuspliner methods for identifying the dominant taxa that contribute to the microbial dynamic differences between control and case groups, with sample size *N* = 20, 30 and the number of time points *T* = 10, 20 in scenarios 1-3, respectively. The proposed MTA method has much higher sensitivity and sufficiently high specificity in all scenarios compared to Permuspliner, although Permuspliner has slightly higher specificity. This indicates that MTA has the ability to identify not only dominant taxa but also those inactive ones. But Permuspliner suffers from limited sensitivity in identifying dominant taxa reliably, especially with *T* = 20, which agrees with its conservative performance in the power section. Thus, the proposed MTA method has better performance in identifying whether or not taxa are dominant than Permuspliner at the taxon level.

**Figure 2:**
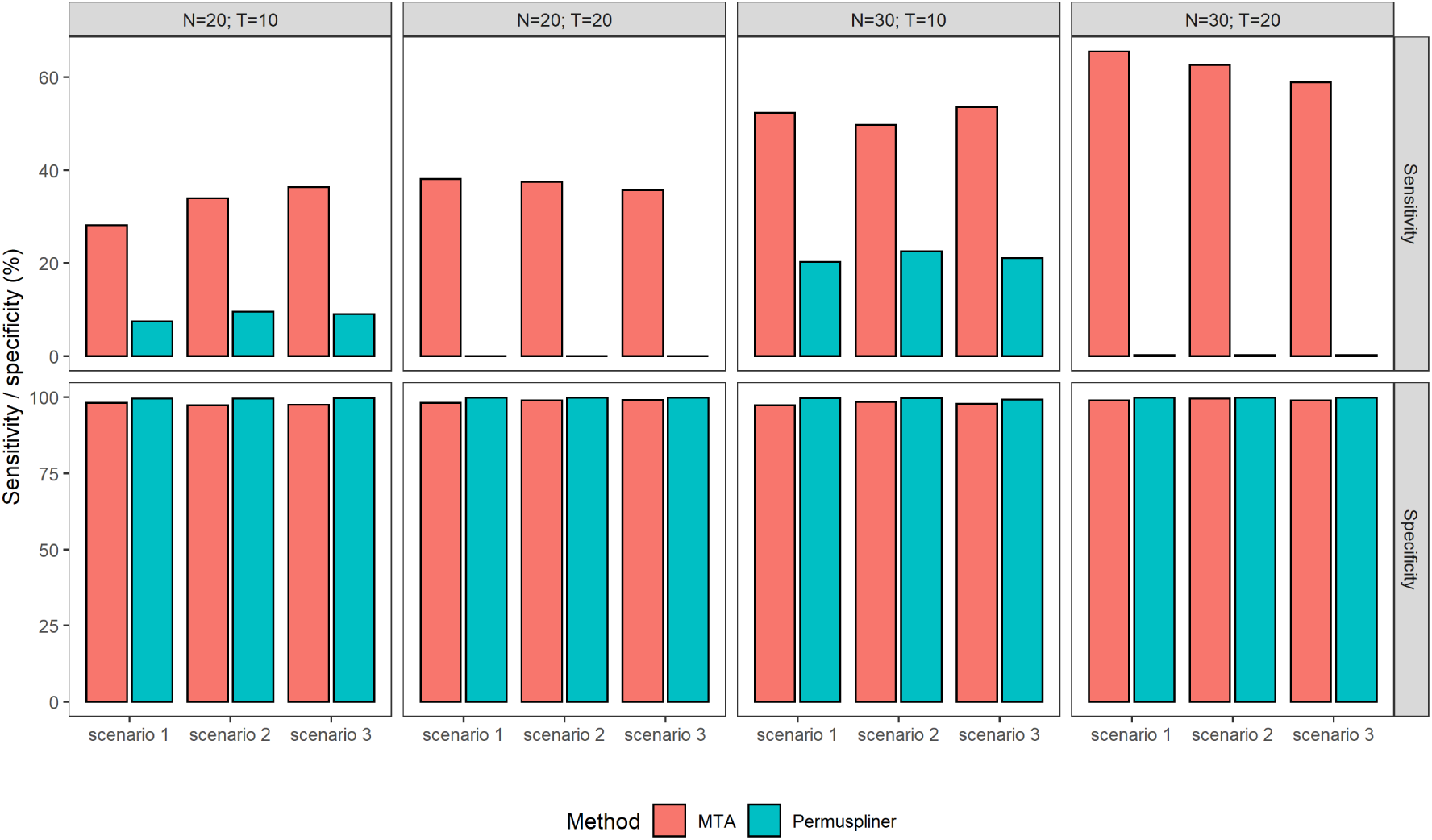
The estimated sensitivity and specificity for identifying the dominant taxa which contribute to the extracted trends with sample size *N* = 20, 30 and the number of time points *T* = 10, 20 in scenarios 1-3, respectively.

Furthermore, the proposed MTA method has the distinct advantage that it not only identifies the dominant taxa that contribute to the extracted common trends, but also quantifies their contributions by the estimated factor scores. This suggests their individualized effects on the difference between two groups and provide additional insights for followup studies. Additional file 2: Figures S1–S3 report the extracted group difference trends and the estimated factor scores for the corresponding dominant taxa in scenarios 1-3 with *N* = 30 and *T* = 10 respectively, produced by our MTA R package.

### Simulation results based on the perturbation method simulation design

To assess the robustness of the proposed MTA method, we further evaluated both methods using the perturbation method simulation design in the original Permuspliner report [21], which is shown in detail in the Additional file 1: Section S3. When the effect directions of the signal taxa at different time points are mixed, the relative performance of MTA and Permuspliner is quite similar to that reported above (the “Simulation results for group comparison” section). MTA exhibits general superior performance in both group and taxon levels evaluation (Additional file 2: Figures S4 and S6). When the effects of the signal taxon across all the time points are in the same direction, which becomes less likely as the number of time points increases, Permuspliner has higher power than MTA, since Permuspliner directly measures the area between the microbial splines of control and case groups for each taxon. However, as the effect size gets larger, the estimated powers of MTA and Permuspliner become similar (Additional file 2: Figure S5). Furthermore, MTA has similar performance with Permuspliner in terms of sensitivity and specificity (Additional file 2: Figure S7). Therefore, considering that the same effect direction across a long period is less likely, the proposed MTA method is robust and has superior performance to Permuspliner in terms of statistical power at the group level, as well as in terms of sensitivity and specificity at the taxon level.

### Simulation results for classification

In this subsection, we evaluate the performance of the proposed classification Algorithm 3 in terms of ROC curve and AUC using the 10-fold CV method based on scenario 2 with different effect sizes. Specifically, we simulated the microbial relative abundances for 300 cases and 300 controls, where 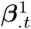 and 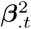 were assigned as below,

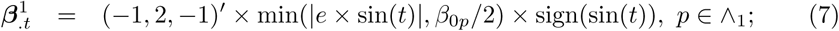

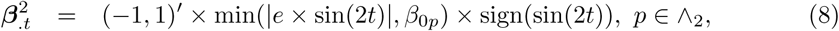

where *e* is the magnitude of effect size and was assigned as 1, 2, 4, 6 and 8, respectively and sign(·) is the sign function. We then randomly divided the 600 subjects into 10 groups of equal size, each of which was in turn used as a testing set to classify a new subject, while the rest were used as a training set to establish the predicted microbial composition matrix for the control and case groups. The ROC curve and AUC were estimated as the average performance over 10 testing sets.

Additional file 2: Figure S8 and Figure 3 present the overall ROC curves and AUCs with effect size *e* = 1, 2, 4, 6, 8, and the number of time points *T* = 10, 20, respectively. As expected, AUC increases as the effect size *e* increases, with the average AUC being 0.63, 0.76, 0.88, 0.95 and 0.96 for *e* = 1, 2, 4, 6, 8 respectively. These results show that the proposed MTA framework has satisfactory performance to classify.

**Figure 3:**
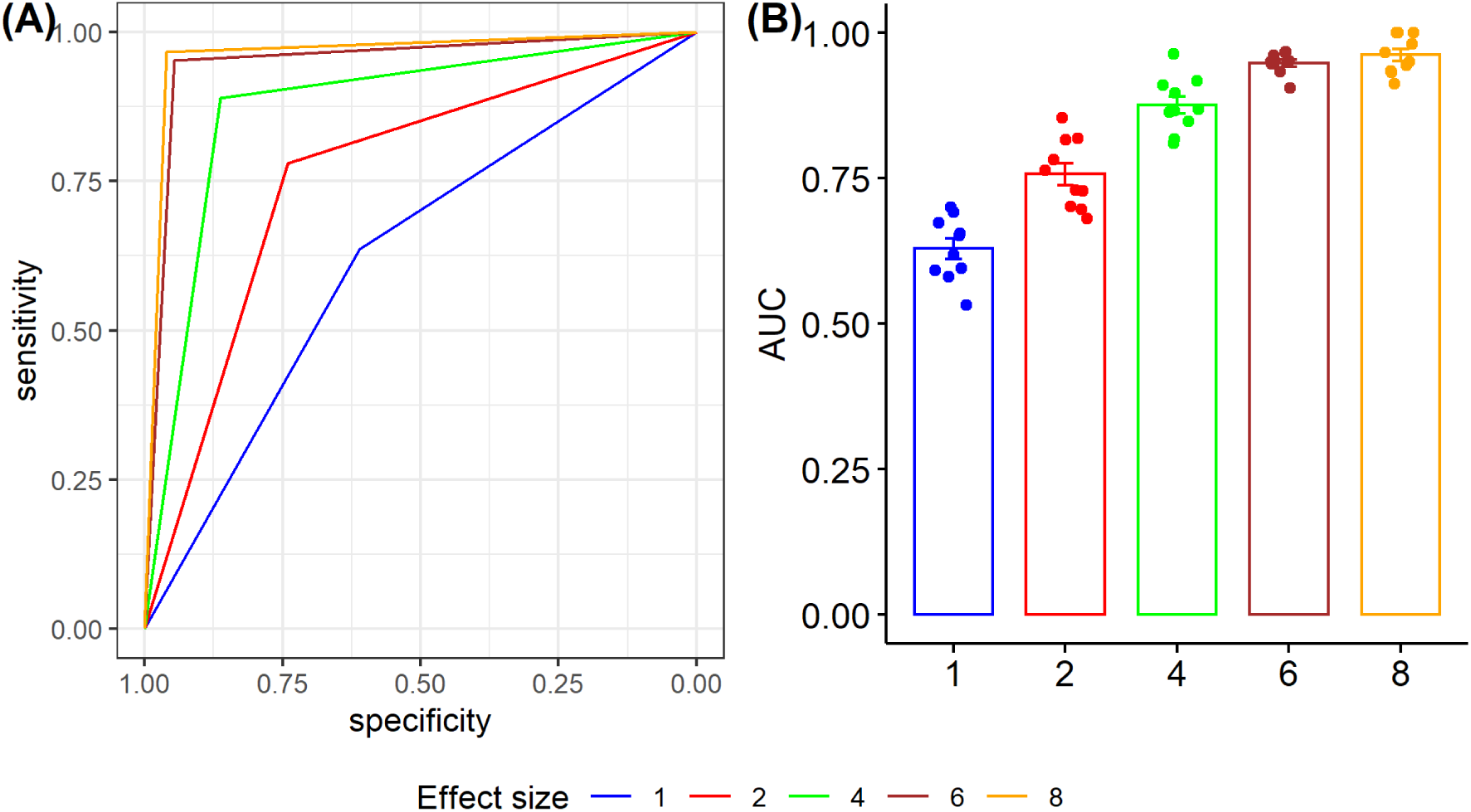
The classification performance of the proposed MTA framework with the number of time points *T* = 20. (A) The overall ROC curves. (B) The mean and standard error of AUCs under various effect sizes based on 10-fold CVs.

## Real data analysis

Livanos et al. (2016) [39] conducted a longitudinal microbiome experiment in mice at risk for developing Type I diabetes (T1D). They found that early exposure to antibiotics altered the gut microbiota and that this shift accelerated T1D onset compared to control mice. Here we apply the proposed MTA method to the longitudinal microbiome data collected in this experiment to examine the changing microbial trend differences between the STAT (antibiotic) and control groups. Specifically, MTA determines whether the overall dynamic patterns present in the STAT and control groups are significantly different or not, identifies which taxa contribute to the identified overall group difference, and subsequently provides the corresponding estimated factor scores for those identified taxa. Algorithm 3 in the MTA framework then is applied to classify the mice based on their longitudinal microbial profiles.

In this study, the fecal samples collected from antibiotic and control mice at 3, 6, 10, and 13 weeks were sequenced to examine their 16S rRNA genes, and their median sequencing depths were 12, 524 ± 3, 295 sequences. The OTU table and taxonomy were determined using the QIIME pipeline [43], and 106 genera were originally observed. As indicated in the “Simulation design” section, we analyzed 35 genera in 17 antibiotic and 20 control male mice at 3, 6, 10 and 13 weeks.

Figure 4(A) reports the primary significant microbial trend determined by the MTA method, with p-value 9.8 × 10^−10^, representing the major difference in the microbial dynamic pattern between antibiotic and control male mice. With MTA, based on the 35 genera, we identify 5 dominant genera (*Lactobacillus, Lachnospiraceae Other, S24-7 Other, Akkermansia*, and *Allobaculum*) that contribute to the microbial trend difference. In Figure 4(B), we report the nonzero estimated factor scores of these 5 dominant genera and they reveal the extent of their contributions to the microbial trend group difference. In contrast, the competing test Permuspliner only identifies 3 significant genera (*Lactobacillus, Coprobacillus, Coriobacteriaceae_Other*) associated with STAT treatment (adjusted p-values < 5%).

**Figure 4:**
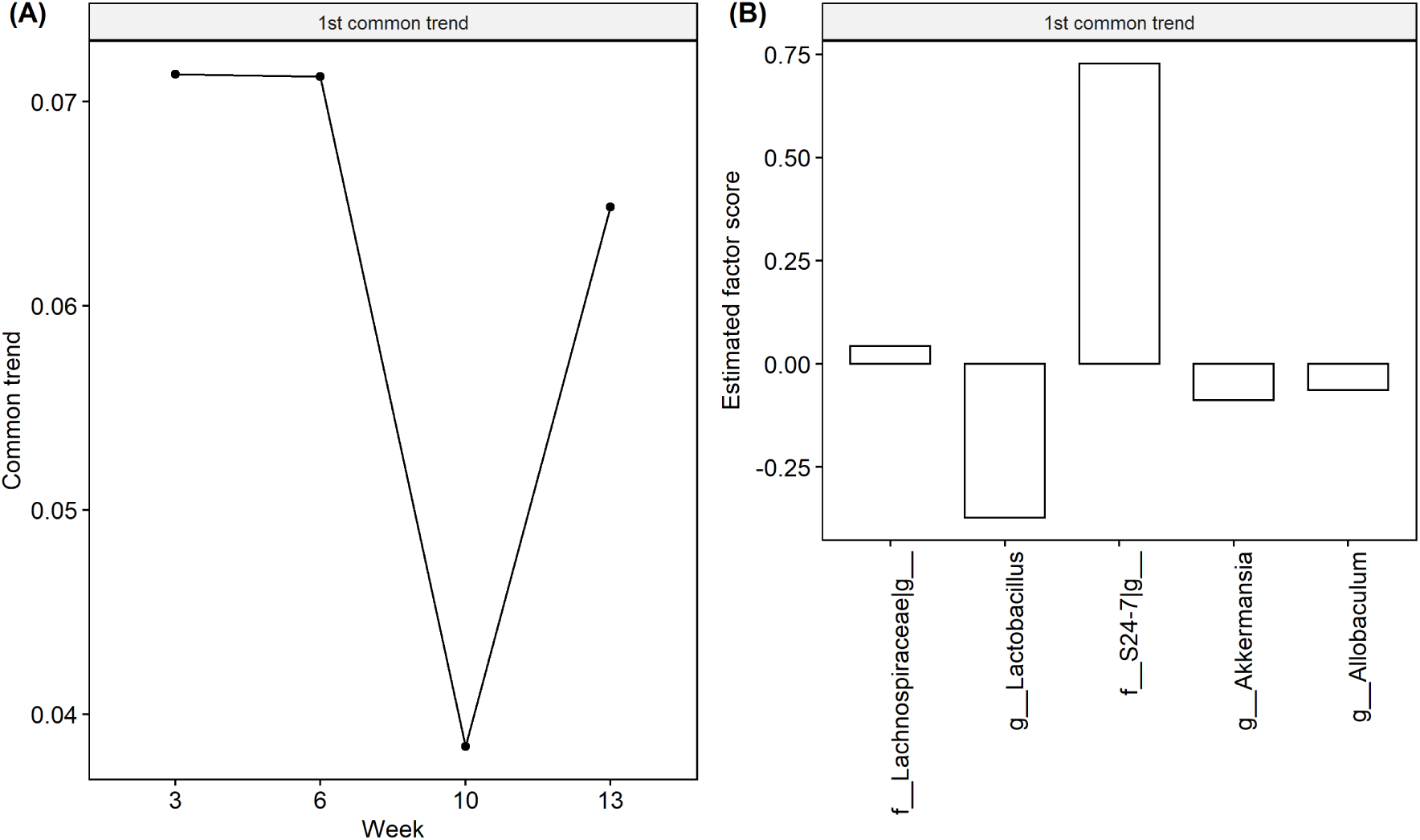
Microbial trend comparison between the STAT and control mice in the real longitudinal murine microbiome data. (A) The microbial trend extracted by MTA represents the significant difference between STAT and control mice. (B) The estimated factor scores for the dominant taxa that contribute to this trend. The number of resamplings *R* = 50.

Figure 5 presents the group average relative abundances over time for each identified genus by either method. Specifically, genus *Lactobacillus* identified by both methods has consistently higher relative abundances at all 4 time points in the control group than in the case group. The genera identified by MTA tend to have different changing directions between the two groups, such as genera *Lachnospiraceae Other, S24-7 Other*, and *Akkermansia*, while Permuspliner fails to identify them, which agrees with the simulation results. The 5 genera identified by MTA are relatively common with their cumulative relative abundances ranging from 46.4% to 97.6% across all 4 time points. In contrast, two of three genera identified by Permuspliner are rarer with their cumulative relative abundances ranging from 0.6% to 31.0% (Additional file 2: Figure S9). Additional file 2: Figure S10 presents a principal coordinate analysis (PCoA) visualization based on the Bray-Curtis dissimilarity index using all 35 original genera; the 5 genera identified by MTA, and the 3 genera identified by Permuspliner at 3, 6, 10 and 13 weeks, respectively. As expected, the PCoA plot based on the genera identified by MTA closely resembles the original plot, while the plot based on the genera identified by Permuspliner presents a completely different pattern. These results illustrate that 5 genera identified by MTA well represent the original microbial diversities among all samples, which demonstrates that MTA is capable of capturing the differential microbial dynamic signals, and representing taxa in the longitudinal microbiome analysis.

**Figure 5:**
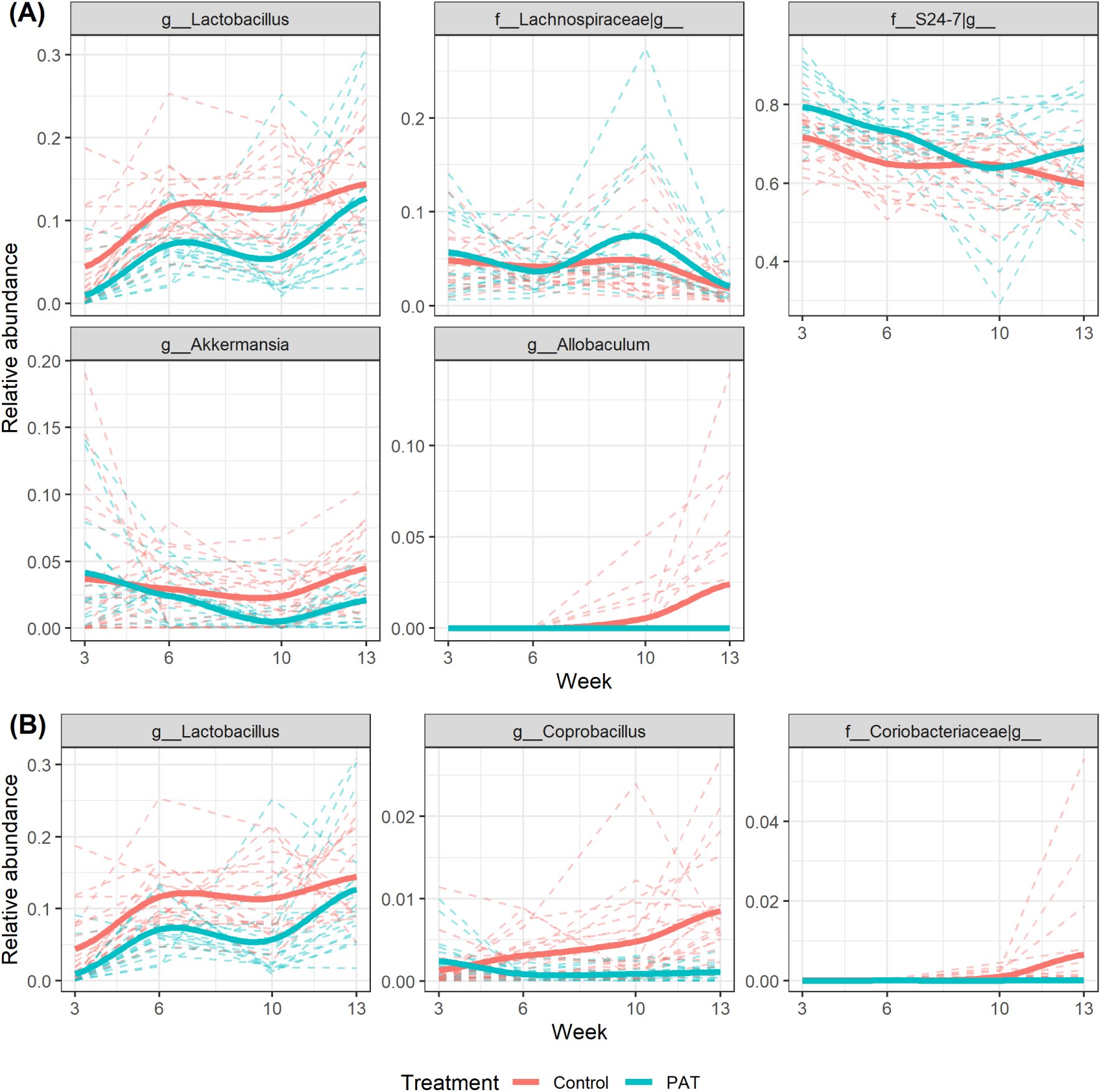
The relative abundances for 7 identified genera at 3, 6, 10 and 13 weeks, respectively. (A) 5 genera identified by the proposed MTA method. (B) 3 genera identified by the Permuspliner method. Genus *Lactobacillus* is identified by both methods.

Figure 6 presents the distances of all mice from the predicted microbial matrices of control and STAT mice which are estimated from the training set based on Algorithm 3. For a given mouse, the training set consists of all other mice. We observe that 18 of 20 control mice (specificity=90.0%) and 13 of 17 antibiotic mice (sensitivity=76.4%) are classified correctly by the proposed distance-based classification algorithm (Algorithm 3), which is superior to the clustering results based on beta diversity measurements at a single time point (Additional file 2: Figure S10) as well as the results in Livanos et al. (2016) [39] based on the hierarchical clustering using the samples at week 6. These results demonstrate the satisfactory performance of the proposed MTA method in classification. Furthermore, the variation of distances shows that STAT mice have more diverse microbial profiles than control mice, which is consistent with their beta diversity measurements shown in Additional file 2: Figure S10.

**Figure 6:**
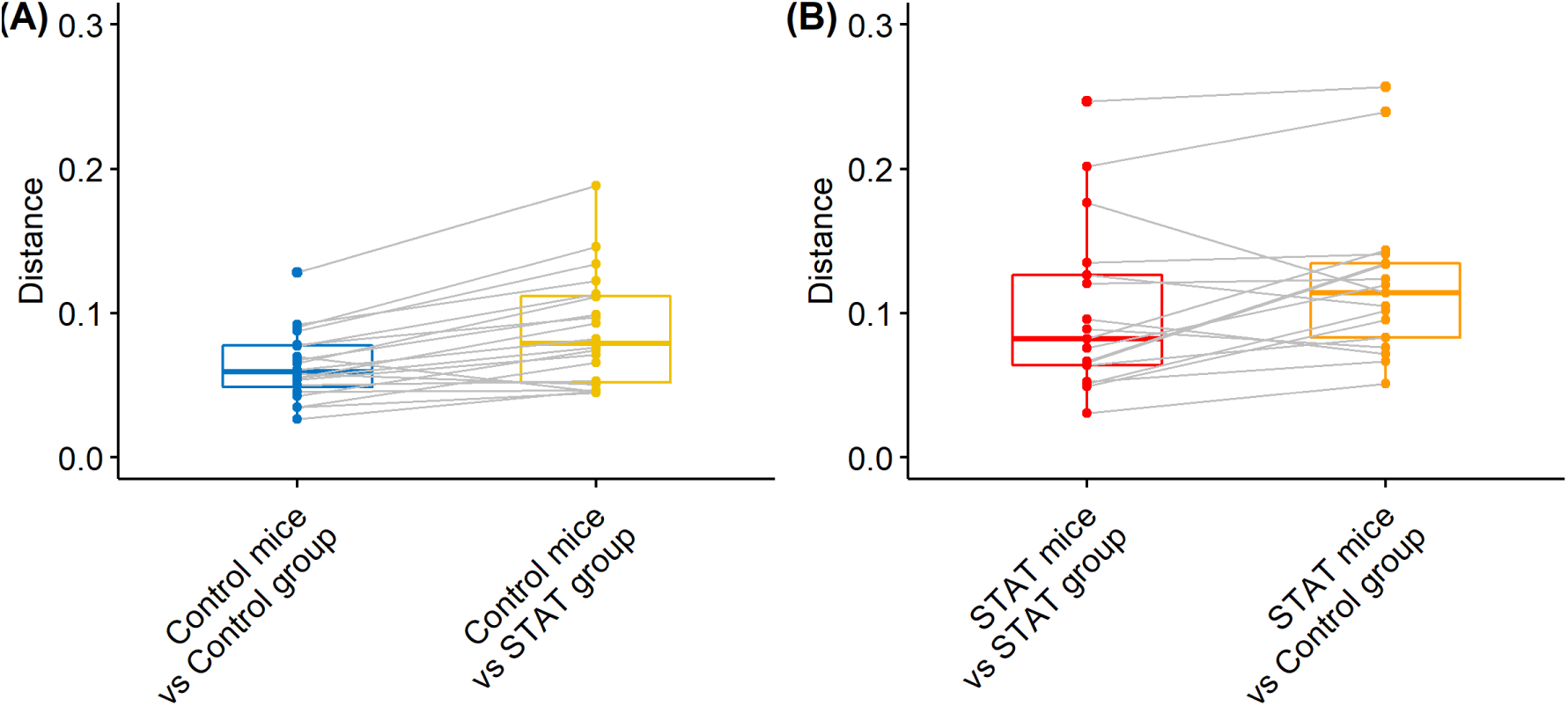
The boxplot of distances calculated based on Algorithm **??**. For a given mouse, the training set involves all other mice except for itself. The distances of its prediction from the predicted microbial matrices of control and STAT mice were separately calculated.

Compared to the significant signals found in Livanos et al. (2016) [39], the proposed MTA method identifies more significant results in terms of dominant taxa identification and mouse classification, which provides insight into the comprehensive characterization of the gut microbiome. This further shows the importance of investigating microbial dynamic changes and their characterization in human diseases based on the longitudinal microbial profiles.

## Discussion

The regularization techniques incorporated in the proposed MTA method provides flexibility to address the distinct concerns of microbiome data. Specifically, MTA employs Lasso penalty, Laplacian penalty, and the smoothing technique to deal with the high dimensionality of taxa, the phylogenetic correlation, and the smoothness of the extracted trends accordingly. The MTA framework assumes that the microbiota acts as an integrated microbial community and that there are a group of key taxa that drive the overall microbial dynamics, rather than assume that each microbe acts alone and needs to be tested individually [2, 44]. This community level integrated analysis enables MTA to provide a systematic view of microbial dynamic responses, which is especially important for understanding complex diseases.

In this paper, MTA analyzes the microbial relative abundance directly, although various log-ratio transformation techniques have been used in dealing with the compositionality of microbiome data [45–47]. Those log-ratio transformations are designed to release the unit sum constraint in the compositional data at a fixed time-point, but using them in the longitudinal microbiome study will alter the inherent microbial trend over time. In Additional file 1: Section S4, we report the results from the additional simulations where we applied the proposed MTA framework to the relative abundances, the corresponding log transformation, additive log-ratio transformation and centered log-ratio transformation, respectively, to illustrate their comparisons under the extensive simulation scenarios. The estimated powers demonstrate that using relative abundances makes MTA capture more dynamic information and produce the highest power in all scenarios. Hence we suggest using the relative abundances directly in the proposed MTA method.

Although this paper focused on binary phenotypes, the proposed MTA framework can be easily generalized for use with more than 2 categories. For group comparisons, Algorithm 2 can be used to compare the paired difference between any two categories and then determine their microbial differential with correction for multiple testing. For the classification, the predicted microbial matrix for each category can be estimated based on the training set, then the distances of a new subject from its microbial profile can be calculated in relation to the predicted microbial matrices of all categories separately, and finally to determine its label by comparing those distances.

## Conclusions

In this paper, we propose a microbial trend analysis (MTA) framework for analyzing the longitudinal microbiome data, which can describe microbial dynamics, differential the group difference, extract key taxa driving the microbial temporal trend, and classify the subjects. Comparing to the competing method Permuspliner, MTA has superior performance in these tasks based on both extensive simulations and real data analyses. Consequently, with the recent proliferation of microbial longitudinal studies, the proposed MTA framework is an attractive analytical tool to study the comprehensive characterization of microbial dynamics and to identify key bacterial species that may affect susceptibility to complex diseases.

## Abbreviations

ALR: additive log-ratio transformation
AUC: area under the curve
BH: Benjamini-Hochberg
CLR: centered log-ratio transformation
CV: cross-validation
IBD: inflammatory bowel disease
ILR: isometric log-ratio transformation
MTA: microbial trend analysis
OTU: operational taxonomic unit
PCoA: principal coordinate analysis
PTA: principal trend analysis
ROC: receiver operating characteristic
STAT: sub-therapeutic antibiotic treatment
T1D: type 1 diabetes.

## Funding

This work was supported in part by National Institutes of Health grants R01DK110014 and U01AI22285, the Fondation Leducq Transatlantic Network, and the Zlinkoff and C&D Funds.

## Availability of data and materials

The longitudinal murine microbiome data [39] we used in our real data analysis is available at the European Bioinformatics Institute database (https://www.ebi.ac.uk) with accession number ERP016357 and Qiita database (https://qiita.ucsd.edu) with accession number 10508. The MTA R package for the proposed microbial trend analysis is publicly available on the web at https://sites.google.com/site/huilinli09/software and https://github.com/chanw0/MTA together with its manual.

## Authors’ contributions

CW developed the microbial trend analysis framework, performed simulation analyses, real data analyses and manuscript writing. JH contributed to real data analyses and manuscript writing. MJB contributed to the biological insights and interpretation, and manuscript writing. HL contributed to the methodological ideas for the proposed framework, simulations, real data analyses, and manuscript writing. All authors read and approved the final manuscript.

## Competing interests

The authors declare that they have no competing interests.

## Consent for publication

Not applicable. All utilized microbiome datasets are publicly available. No consent for publication was required for this study.

## Ethics approval and consent to participate

All utilized microbiome datasets are publicly available. No ethics approval or consent to participate was required for this study.

## Figures

### Additional Files

#### Additional file 1

**Section S1:** Laplacian matrix construction.

**Section S2:** The derivation of Algorithm 1.

**Section S3:** Simulation study based on the perturbation method simulation design.

**Section S4:** Comparison between using relative abundance and using transformed relative abundances.

#### Additional file 2

**Figure S1.** The proposed MTA method for the comparison between case and control groups in scenario 1 with sample size *N* = 30 and the number of time points *T* = 10. (A) The microbial trend extracted by MTA represents the significant difference between case and control groups. (B) The average and standard error of estimated factor scores for the dominant taxa that contribute to the extracted trend respectively.

**Figure S2.** The proposed MTA method for the comparison between case and control groups in scenario 2 with sample size *N* = 30 and the number of time points *T* = 10. (A) Two microbial trends extracted by MTA represent the significant difference between case and control groups. (B) The average and standard error of estimated factor scores for the dominant taxa that contribute to those two trends respectively.

**Figure S3.** The proposed MTA method for the comparison between case and control groups in scenario 3 with sample size *N* = 30 and the number of time points *T* = 10. (A) The microbial trend extracted by MTA represents the significant difference between case and control groups. (B) The average and standard error of estimated factor scores for the dominant taxa that contribute to the extracted trend respectively.

**Figure S4.** Empirical power for testing the difference between case and control groups with sample size *N* = 20, 30 and the number of time points *T* = 10, 20 under 1X and 2X magnitudes of perturbation, respectively. Here ***z*** = (0, 0.3, 0.45, 0.2, −0.3, 0.3, −0.3, −0.2, −0.1, 0)′ and ***z*** = (0, 0.2, 0.5, 0.3, 0.2, −0.2, −0.4, −0.4, −0.2, 0.2, 0.5, 0.2, −0.3, −0.4, −0.2, 0.2, 0.4, 0.2, 0.1, 0)′ for *T* = 10, and 20, respectively.

**Figure S5.** Empirical power for testing the difference between case and control groups with sample size *N* = 20, 30 and the number of time points *T* = 10, 20 under 1X and 2X magnitudes of perturbation, respectively. Here ***z*** = (0, 0.1, 0.2, 0.3, 0.4, 0.3, 0.2, 0.1, 0.1, 0)′ and ***z*** = (0, 0.05, 0.1, 0.2, 0.2, 0.2, 0.3, 0.3, 0.2, 0.2, 0.2, 0.2, 0.3, 0.3, 0.2, 0.2, 0.2, 0.2, 0.1, 0)′ for *T* = 10, and 20, respectively.

**Figure S6.** The estimated sensitivity and specificity for identifying dominant taxa which contribute to the extracted trends with sample size *N* = 20, 30 and the number of time points *T* = 10, 20 under 1X and 2X magnitudes of perturbation, respectively. Here ***z*** = (0, 0.3, 0.45, 0.2, −0.3, 0.3, −0.3, −0.2, −0.1, 0)′ and ***z*** = (0, 0.2, 0.5, 0.3, 0.2, −0.2, −0.4, −0.4, −0.2, 0.2, 0.5, 0.2, −0.3, −0.4, −0.2, 0.2, 0.4, 0.2, 0.1, 0)′ for *T* = 10, and 20, respectively.

**Figure S7.** The estimated sensitivity and specificity for identifying dominant taxa which contribute to the extracted trends with sample size *N* = 20, 30 and the number of time points *T* = 10, 20 under 1X and 2X magnitudes of perturbation, respectively. Here ***z*** = (0, 0.1, 0.2, 0.3, 0.4, 0.3, 0.2, 0.1, 0.1, 0)′ and ***z*** = (0, 0.05, 0.1, 0.2, 0.2, 0.2, 0.3, 0.3, 0.2, 0.2, 0.2, 0.2, 0.3, 0.3, 0.2, 0.2, 0.2, 0.2, 0.1, 0)′ for *T* = 10, and 20, respectively.

**Figure S8.** The classification performance of the proposed MTA framework with the number of time points *T* = 10. (A) The overall ROC curves. (B) The mean and standard error of AUCs under various effect sizes based on 10-fold CVs.

**Figure S9.** The relative abundances for the genera identified by MTA (5 genera) and Permuspliner (3 genera) at 3, 6, 10 and 13 weeks respectively.

**Figure S10.** Beta diversity analysis for all male mice based on the Bray-Curtis dissimilarity index. The PCoA is evaluated based on: (A1)-(A4) 35 original genera; (B1)-(B4) 5 genera identified by the proposed MTA method; and (C1)-(C4) 3 genera identified by the competing method Permuspliner at 3, 6, 10 and 13 weeks respectively. Points represent samples.

**Figure S11.** Estimated powers (%) of the proposed MTA method for testing the microbial trend difference between case and control groups using different transformed relative abundances, with sample size *N* = 20 and the number of time points *T* = 10 respectively. Specifically, the proposed MTA method was applied on the relative abundances directly (None; red bar), the log transformation (Log; blue bar), ALR (additive log-ratio transformation; darkgreen bar), and CLR (centered log-ratio transformation; orange bar) respectively under the scenarios 1-3, which were constructed as in the “Simulation design” section.

**Figure S12.** Estimated powers (%) of the proposed MTA method for testing the microbial trend difference between case and control groups using different transformed relative abundances, with sample size *N* = 20 and the number of time points *T* = 10 respectively. Specifically, the proposed MTA method was applied on the relative abundances directly (None; red bar), the log transformation (Log; blue bar), ALR (additive log-ratio transformation; darkgreen bar), and CLR (centered log-ratio transformation; orange bar) respectively under the scenarios 1-3, which were constructed as in the Additional file 1: Section S4.

